# Non-canonical activation of the ER stress sensor ATF6 by *Legionella pneumophila* effectors

**DOI:** 10.1101/2020.06.26.174185

**Authors:** Nnejiuwa U. Ibe, Shaeri Mukherjee

## Abstract

The intracellular bacterial pathogen *Legionella pneumophila* (*L.p.*) secretes ~330 effector proteins into the host cell to sculpt an Endoplasmic Reticulum (ER)-derived replicative niche. We previously reported five *L.p.* effectors that inhibit IRE1, a key sensor of the homeostatic unfolded protein response (UPR) pathway. In this study, we discovered a subset of *L.p.* toxins that selectively activate the UPR sensor ATF6, resulting in its cleavage, nuclear translocation and target gene transcription without affecting other UPR sensors such as PERK. In a deviation from the conventional model, this *L.p*. dependent activation of ATF6 does not require its transport to the Golgi or its cleavage by the S1P/S2P proteases. We believe that our findings highlight the unique regulatory control that *L.p*. exerts upon the three UPR sensors and expand the repertoire of bacterial proteins that selectively perturb host homeostatic pathways.

## Introduction

Several intracellular pathogens, including *Legionella pneumophila (L.p.)*, expertly manipulate host cell function to create their replicative niche. *L.p.* uses the specialized Dot/Icm Type IVB secretion system (T4SS) to translocate roughly 300 bacterial effector proteins into the host cytosol (Berger & Isberg, 1993, Hubber & Roy, 2010, Isberg, O’Connor et al., 2009, Vogel, Andrews et al., 1998). Once deposited into the cytosol, the effectors target a vast array of host proteins and can influence diverse biological processes which permit the use of *L.p*. as a tool to uncover novel biological mechanisms. During infection, *L.p.* uses its effectors to prevent fusion of the *Legionella*-containing vacuole (LCV) with the host endosomal machinery. Instead these effectors facilitate the remodeling of the LCV into a compartment that supports pathogen replication (Marra, Blander et al., 1992, Roy, Berger et al., 1998, Wiater, Dunn et al., 1998). Though fusion with lysosomes is evaded during infection, there is substantial interaction between the LCV and other host organelles including the endoplasmic reticulum (ER) (Horwitz & Silverstein, 1983, Swanson & Isberg, 1995, Tilney, Harb et al., 2001).

The ER-LCV interactions take on different forms as LCV maturation progresses. *L.p.* induces tubular ER rearrangements and intercepts ER-derived vesicles destined for the Golgi early in infection (Kotewicz, 2017). However, the mature LCV is studded with ribosomes and reticular ER proteins presenting a vacuolar environment that is substantially different and highlights the complexity of interactions between the ER and LCV. The ER serves as a critical regulatory site for protein and membrane lipid biosynthesis, and imbalances in protein load or membrane lipid perturbations can disrupt many of its vital homeostatic functions (Rapoport, 2007, Halbleib, 2017). The unfolded protein response (UPR) serves as a prominent regulatory pathway in the ER that has been shown to respond to the burden of accumulating unfolded or misfolded proteins in the ER (Ron & Walter, 2007). In mammalian cells, the UPR is coordinated by three ER-localized transmembrane proteins, inositol-requiring protein-1 (IRE1), protein kinase RNA (PKR)-like ER kinase (PERK) and activating transcription factor-6 (ATF6), each of which initiate pathways designed to modulate the cellular response (Cox, Shamu et al., 1993, Lee, Tirasophon et al., 2002, Mori, Ma et al., 1993).

ATF6 is a type II transmembrane protein that is retained in the ER under normal homeostatic conditions through interactions with the resident chaperone BiP/GRP78 (Shen, Chen et al., 2002). Upon accumulation of unfolded proteins, the ER stress stimulates ATF6 translocation from the ER to the Golgi. At the Golgi, ATF6 is sequentially cleaved first by site-1 protease (S1P) in the luminal domain, then by site-2 protease (S2P) liberating the cytosolic ATF6-N terminal fragment, ATF6(NT) (Shen & Prywes, 2004, Ye, Rawson et al., 2000). Once cleaved, ATF6(NT) is recruited to the nucleus where it binds to cis-acting ER stress response elements (ERSE) in the promoter region of UPR target genes (Kokame, Kato et al., 2001, Yoshida, Okada et al., 2000). ATF6 activation is thought to facilitate cytoprotective adaptation to ER stress through the regulation of genes that improve protein folding and processing in the ER. ATF6 has been shown to suppress the UPR induced apoptotic program once the stress is resolved (Wu et al., 2007), highlighting the pro-survival contributions of this signaling network.

Studies emphasizing cross talk between the UPR and bacterial infection have revealed an interconnectedness of ER stress sensing and pathogen sensing mechanisms in the cell (Li, Wang et al., 2011, Urano, Wang et al., 2000). Pathogenic perturbations endured during infection can impact ER homeostasis in a manner that can also induce ER stress responses. Intracellular pathogens across all kingdoms, from virus to protozoans, have devised strategies to subvert or utilize one or more UPR program to benefit survival and replication within the host (Celli & Tsolis, 2015, Galluzzi, Diotallevi et al., 2017, Verchot, 2016). As further evidence, studies have demonstrated pathogen-mediated targeting of ATF6 can be beneficial for survival (Ambrose & Mackenzie, 2013, Hou, Dong et al., 2019, Hou, Wei et al., 2017) and replication (Yoshikawa, Sugimoto et al., 2020). Our previous analysis on *L.p.* mediated manipulation of the UPR revealed a dynamic reduction in full length ATF6 protein levels during infection (Treacy-Abarca & Mukherjee, 2015). To further understand the relationship between *L.p.* infection and ATF6 processing, we sought to understand the mechanism by which *L.p.* modulates the ATF6 pathway. Here, we present evidence of a unique, non-canonical mode of ATF6 activation by *L.p.* that does not rely on host proteins, that were previously thought to be essential for ATF6 activation. Surprisingly, we discover novel *L.p.* effectors that play a role in activation of ATF6 during infection.

## Results

### Proteasome-dependent degradation pathways are not required for ATF6 loss during L.p. infection

Upon infection with wild type *L.p.* (*WT-L.p.*), we observed near complete processing of endogenous full length ATF6 (Figure 1A). The level of processing was similar to that induced by the strong reducing agent and non-specific ER stress inducer, dithiothreitol (DTT) (Figure 1A). An isogenic strain of *L.p.* that lacks a functional secretion system (Δ*dotA-L.p.*) was unable to downregulate ATF6 protein levels, suggesting that one or more bacterial effectors could be responsible for its targeting (Figure 1A). Processing of ATF6 through regulated intramembrane proteolysis (RIP) by the S1P and S2P proteases has been studied extensively (Okada, 2003; Ye, 2000), yet degradative processing events have also been shown to control ATF6 levels even in the absence of ER stress (Hong, Li et al., 2004, Horimoto, Ninagawa et al., 2013). Interestingly, protein synthesis attenuation during *L.p.* infection has been shown to influence the IRE-1 branch of the UPR (Hempstead, 2015, Treacy-Abarca, 2015); but its impact on ATF6 has not been elucidated.

**Figure 1.**
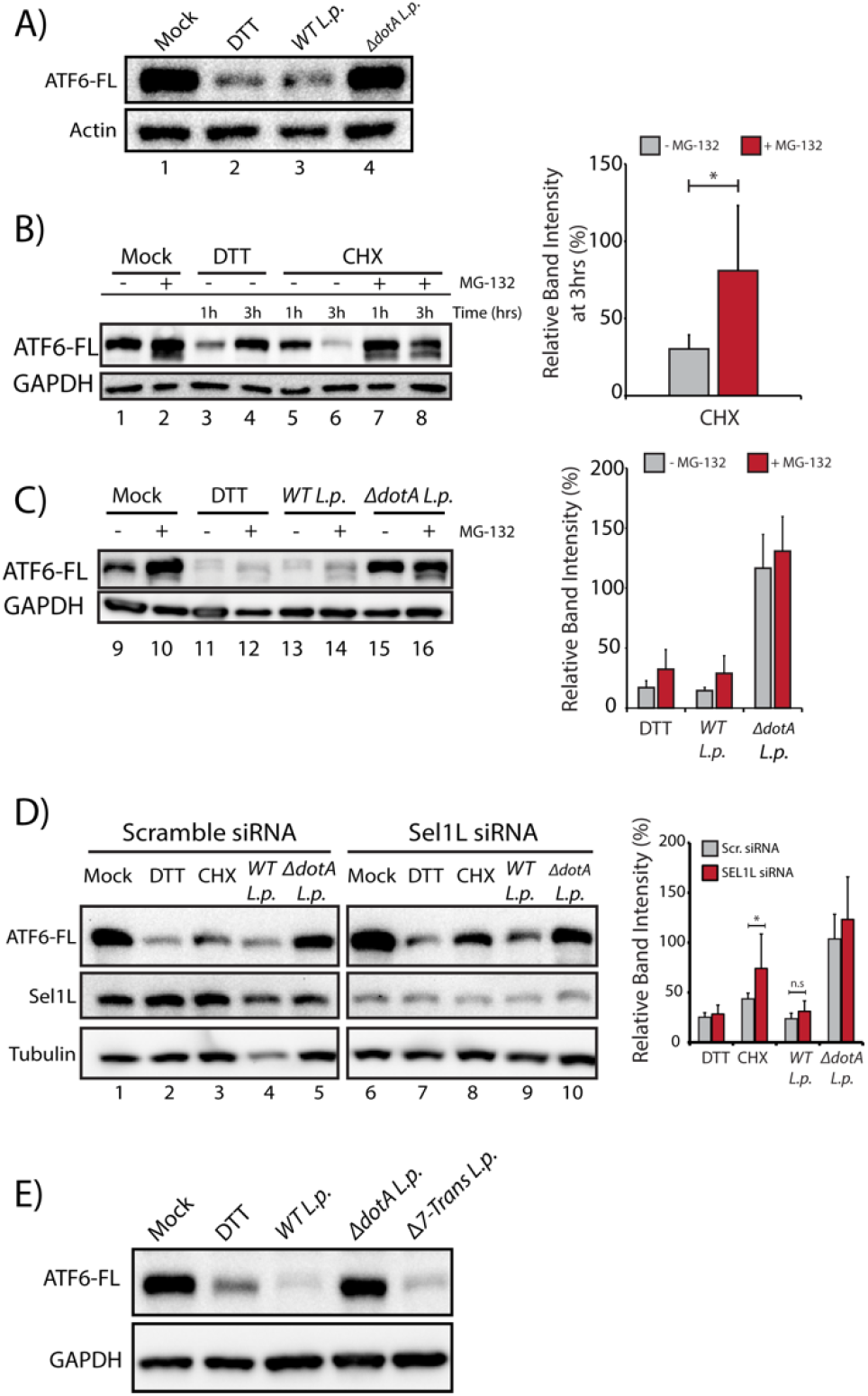
*L.p.* infection induces ATF6 processing independently of proteasomal degradation and ERAD. (A) HEK293-FcγR cells were treated with 1 mM DTT for 1 hour or infected with *Legionella* pneumophila (*WT* or *ΔdotA*) for 6 hours prior to harvesting. Lysates were subjected to immunoblot analysis using anti-ATF6 and anti-Actin antibody. The precursor full-length ATF6 (ATF6-FL) fragment is shown. (B and C) HEK293-FcγR cells were pre-treated with 20 uM MG-132 for 3 hours prior to indicated treatment conditions. (B) Cells were treated with 1mM DTT and 25 μM Cycloheximide (CHX) for 1 hour or 3 hours as indicated, or (C) infected with *WT*- or Δ*dotA-L.p.* for 6 hours (MOI = 5) and analyzed by immunoblot using anti-ATF6 and anti-GAPDH antibody. Quantitation from full length ATF6 signal from replicate experiments (n=3). (D) Cell lysates from scramble or *SEL1L* siRNA transfected HEK293-FcγR cells were analyzed by immunoblot using anti-ATF6, anti-Sel1L, and anti-Tubulin antibodies. CHX treatment was performed with 25 μM CHX for 2 hours. Quantitation from full length ATF6 signal from replicate experiments (n=3). (E) HEK293-FcγR cells were treated with 1 mM DTT or infected with *WT L.p.*, *ΔdotA L.p.* and *Δ7-translation* (*Δ7-Trans) L.p.* strain as described in (A).

To test if proteasomal degradation contributed to the observed loss of ATF6, we monitored ATF6 processing in the presence or absence of proteasome inhibition, under conditions of protein synthesis arrest, or during *L.p.* infection. HEK293 cells stably expressing the Fcγ receptor (HEK293-FcγR), to allow for antibody-mediated opsonization of *L.p.*, were pre-treated for 3 hours with the proteasome inhibitor MG-132 or control media. ER stress induction using DTT led to rapid ATF6 processing after 1 hour, whereas prolonged exposure to DTT for 3 hours resulted in recovery of ATF6 signal due to autoregulatory feedback from UPR induction (Figure 1B). In contrast to UPR induction, cells treated with protein synthesis inhibitor cycloheximide (CHX) showed loss of full length ATF6 signal after 3 hours post-treatment (Figure 1B). While CHX treatment alone resulted in reduced levels of ATF6, pre-treatment with MG-132 stabilized ATF6 in the presence of CHX to pre-treatment levels (Figure 1B), consistent with previously described observations (Haze et al., 1999). We next tested whether proteasomal inhibition could stabilize ATF6 protein levels in cells infected with *WT-L.p.* or *ΔdotA-L.p.* strains (Figure 1C). Similar to UPR induction using DTT (Figure 1), MG-132 treatment did not protect ATF6 from processing during WT *L.p.* infection (Figure 1C). When cells were infected *ΔdotA L.p.*, ATF6 remained at pre-infection levels and treatment with MG-132 did not have a significant impact (Figure 1C). Previous studies have identified ATF6 as an ER-associated degradation (ERAD) substrate that undergoes constitutive degradation mediated by SEL1L (Horimoto et al., 2013). It was shown that ATF6 is a short-lived protein with a half-life less than 2 hours and the stability of ATF6 can be markedly increased by *Sel1L* knock-out (Horimoto et al., 2013). To test if Sel1L dependent ERAD contributes to loss of ATF6 during infection, we next compared ATF6 processing in HEK293-FcγR cells that were treated with non-targeting or *SEL1L*-targeting siRNA. SEL1L knock-down led to an increase in ATF6(FL) levels by 1.5-fold in samples not treated with CHX (Figure 1D). Similarly, while CHX treatment for 2 hours caused a reduction in ATF6(FL) levels in non-targeting siRNA cells, ATF6(FL) levels were again increased by 1.5-fold in *SEL1L* knock-down cells (Figure 1D). However, there was no significant increase in ATF6(FL) signal intensity under *L.p.* infection when normalized to loading controls (Figure 1D). As *L.p.* infection causes downregulation of protein synthesis (Belyi, Niggeweg et al., 2006, Fontana, Banga et al., 2011, Tzivelekidis, Jank et al., 2011), we considered this deregulation might contribute to loss of ATF6 during infection. To evaluate the impact of protein synthesis inhibition, we tested ATF6 processing in a *L.p.* strain lacking T4SS effectors that are known to block protein synthesis; Δ7-Translation *(Δ7-Trans-L.p.*) *(Barry, Fontana et al., 2013, Fontana et al., 2011)*. Consistent with our previous results, we observed ATF6 processing to similar levels under infection using the *Δ7-Trans-L.p.* strain as seen with *WT-L.p.* (Figure 1E). Together, these experiments suggest ATF6 processing during *L.p.* infection is not a direct result of enhanced proteasomal degradation or a consequence of protein synthesis arrest.

### *L. p. induces ATF6* mediated gene induction

We next considered how ATF6 processing affected its distal function as a nuclear transcription factor. To evaluate transcriptional activation, we used RT-qPCR to compare mRNA levels of ATF6 target genes, including UPR regulator/ER chaperone *BiP (HSPA5)*, in HEK293-FcγR cells under conditions of ER stress and *L.p.* infection. UPR induction with DTT increased expression of ER quality control genes *BiP* and *HERPUD1* by greater than 5-fold compared to control (DMSO-treated) cells (Figure 2A). Interestingly, these ATF6-regulated genes were also induced in wild-type infected cells by greater than five-fold. When compared to protein synthesis inhibition using CHX, we found CHX did not induce *BiP* gene expression (Figure 2B). To gain more insight into ATF6 induction patterns during *L.p.* infection, we examined the gene activation profile of *BiP* over the course of infection. Analysis of *BiP* mRNA expression using RT-qPCR indicated a spike in *BiP* expression between 4-5 hours post DTT treatment (Figure 2C). When the expression profile was examined under avirulent *ΔdotA*-*L.p.* infections, *BiP* expression spiked between 1-3 hours post-infection, but quickly dropped to pre-infection levels by 5 hours post-infection. In contrast, infection with wild-type *L.p.* stimulated *BiP* expression around 3 hours post-infection, with pronounced gene induction even after 5 hours post-infection. The results were corroborated when ATF6 processing was monitored over the same time period. As shown previously, ATF6 is rapidly cleaved under DTT treatment (30 minutes), but the ATF6(FL) signal recovers after sustained stress due to autoregulatory feedback (Figure 2D). In comparison, *ΔdotA*-*L.p.* infection did not induce significant changes in ATF6(FL) levels over the course of infection (Figure 2D). Yet, the appearance of a cleavage product at 1 hour correlated with the spike in *BiP* mRNA expression (Figure 2D). When monitoring ATF6 processing under wild-type *L.p.* infection, the ATF6 processing profile also correlated with the changes in *BiP* mRNA expression. ATF6 signal intensity at 1-hour post-infection was approximately 90% of pre-infection levels (Figure 2E). Within 3 hours of infection, the ATF6 signal intensity had decreased by roughly 50%. By 5 hours post-infection, *BiP* mRNA levels peaked and the ATF6 signal intensity dropped below 25% of pre-infection levels. In contrast to pharmacological ER stress induction, the loss of ATF6(FL) signal persisted throughout the infection. Taken together, the gene expression changes and ATF6 processing analysis demonstrate activation of the ATF6 pathway during *L.p.* infection. It is possible that *L.p.* could induce ATF6 downstream gene activation via effector(s) that don’t require ATF6 cleavage. We thus determined whether ATF6 itself was required for UPR gene induction during infection. The ATF6 gene was targeted for knockdown in HEK293-FcγR cells using siRNA achieving greater than 80% knockdown efficiency (Figure 2F). Though ATF6 knockdown nearly ablated *BiP* induction using DTT, ATF6 knockdown under *L.p.* infection markedly reduced *BiP* mRNA induction, suggesting that the endogenous ATF6 cleavage is indeed being utilized to induce downstream gene activation. However, *BiP* mRNA levels were still elevated nearly 4-fold compared to chemical induction in the knockdown cells (Figure 2G). The residual *BiP* induction in *WT-L.p*. infection could be the result of the remnant ATF6 produced from incomplete knockdown. An alternate scenario is that certain *L.p.* effectors might bypass the requirement for ATF6 cleavage by directly inducing ATF6 gene activation, suggestive of a redundant strategy often employed by *L.p*.. Taken together, the data suggest that *L.p.* infection stimulates ATF6 processing and targets gene induction.

**Figure 2.**
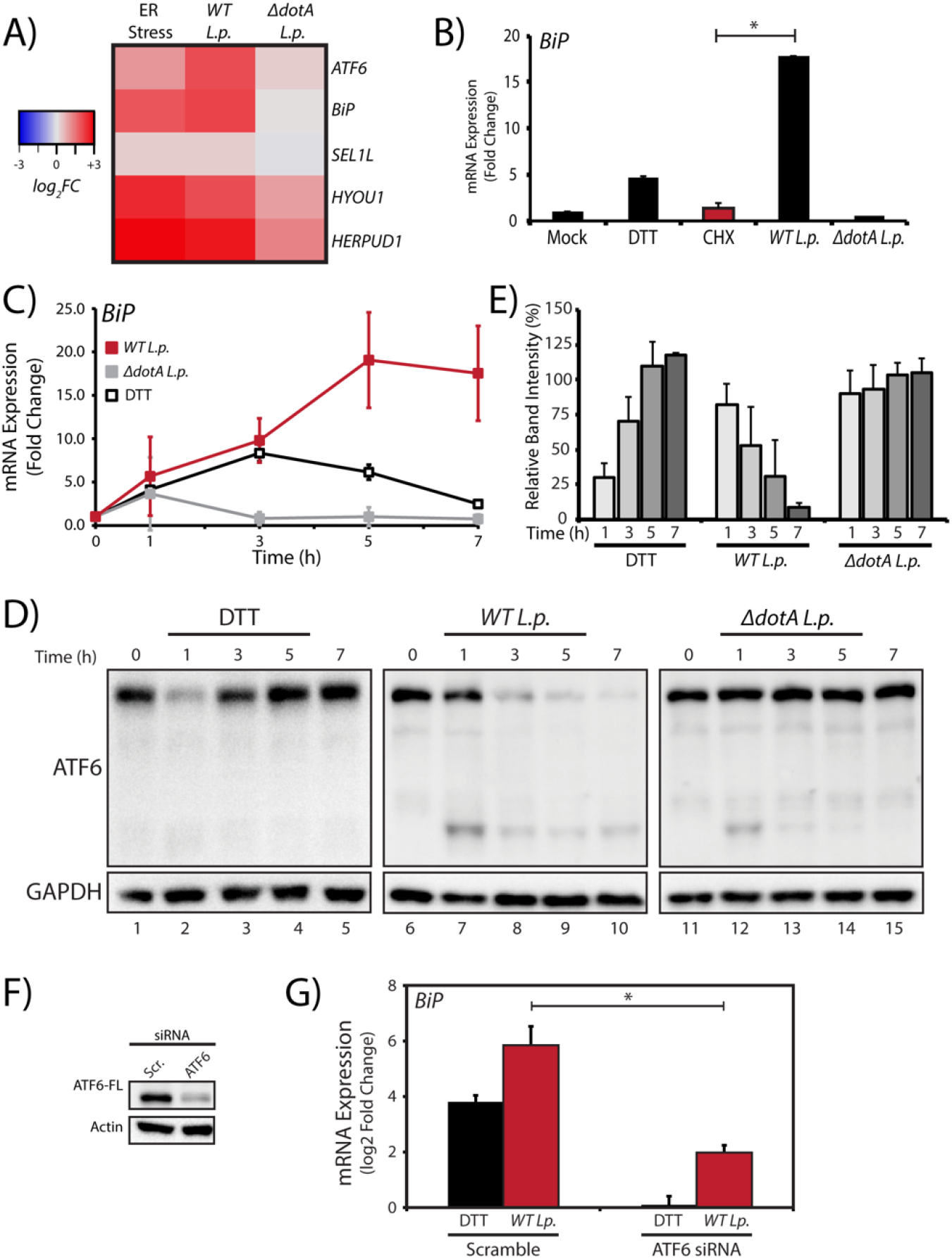
*L.p.* infection stimulates ATF6 target gene induction. (A) HEK293-FcγR cells were treated with 1 mM DTT for 6 hours or infected with *L.p.* (*WT* or *ΔdotA*) for 6 hours prior to harvesting and RT-qPCR analysis using ER quality control genes *ATF6*, *BiP*, *SEL1L*, *HYOU1*, *HERPUD1*. Fold change shown relative to *GAPDH*. (B) Cells were treated with 1mM DTT or 25 μM CHX for 6 hours, or infected with *L.p.* (*WT* or *ΔdotA*) for 6 hours. RT-qPCR analysis of *bip* was performed and fold change was calculated against GAPDH as a reference gene. (C, D) *L.p.* infected HEK293-FcγR cells were harvested post-infection at indicated times, then (C) analyzed by RT-qPCR using primers against *bip,* or (D) lysed and analyzed by immunoblot using anti-ATF6 and anti-GAPDH antibodies. MOI = 100. (E) Quantitation from full length ATF6 signal from replicate experiments relative time point 0 (n=3). (F, G) *ATF6* siRNA or scramble siRNA transfected HEK293-FcγR cells were analyzed by (F) immunoblot using anti-ATF6 and anti-actin antibodies, or (G) analyzed by RT-qPCR using primers against *BiP*.

### *L. p.* induces a non-canonical ATF6 activation

*L.p.’s* vast effector repertoire permits its subversion of numerous host pathways (Alix, 2011, Cornejo, 2017, Noack, 2020). However, it remains unknown which aspects of ATF6 activation during infection are host-driven or effector-driven. To further understand the mechanism of ATF6 activation, we performed pharmacological inhibition of host pathways that are thought to be essential for ATF6 activation. First, we utilized Ceapin inhibitors that selectively block ATF6 activation by inhibiting its translocation to the Golgi (Gallagher, Garri et al., 2016, Gallagher & Walter, 2016). Treatment of cells with DTT resulted in processing of ATF6 (Figure 3A) and a greater than 5-fold induction of *BiP* mRNA (Figure 3B). In contrast, co-treatment of cells with DTT and Ceapin A7 (+A7) partially protected ATF6 from cleavage (Figure 3A) and reduced gene activation of *BiP* by nearly 50%. As shown earlier, *WT-L.p.* infection alone for 6 hours leads to complete processing of ATF6 and strong induction of *BiP* mRNA. Yet surprisingly, pre-treatment with Ceapin A7 did not prevent processing of ATF6 by the *WT-L.p.* strain (Figure 3A and 3B). Additionally, *BiP* mRNA persisted at levels similar to untreated infections. This result suggests that ATF6 translocation from the ER to the Golgi is not a pre-requisite step required for its processing during infection. *L.p.* is known to exploit ER-to-Golgi trafficking and individual effectors have been identified that can disrupt Golgi homeostasis (Mukherjee, Liu et al., 2011). As disrupted homeostasis could result in mis-localization of Golgi proteins, the impact of direct inhibition of S1P on ATF6 activation was tested using the inhibitor PF-429242 (Lebeau, Byun et al., 2018). Inhibition of S1P proteolysis activity using PF-429242 (PF) in the presence of DTT resulted in the appearance of a slower migrating species of ATF6 at a higher molecular weight likely due to extensive glycosylation in the Golgi (Figure 3A). Further validating S1P inhibition, *BiP* mRNA induction was reduced by nearly 80% compared to DTT treatment alone (Figure 3B). Remarkably, S1P inhibition did not alter ATF6 processing in *WT-L.p.* infected cells, and *BiP* mRNA was induced to similar levels as in untreated infected cells (Figure 3A and Figure 3B). The cleavage of ATF6 during *L.p.* infection even in the presence of Ceapin A7 and S1P inhibition are suggestive of an alternative proteolytic mechanism induced by *L.p.*. Because the S1P protease was not required for *L.p.* induced ATF6 processing, we next tested whether the canonical cleavage sites were a prerequisite for cleavage. To test this, a construct harboring a point mutation at the ATF6 S1P cleavage site ATF6(R415A/R416A) was used (Figure 3C). HEK293-FcγR cells transiently expressing GFP-ATF6(R415A/R416A) were treated with UPR inducer DTT or infected with *WT*-*L.p*. or *ΔdotA*-*L.p*. strains. DTT treatment indicated processing of full-length ATF6 was greatly impaired in the S1P cleavage site mutant construct compared to wild type ATF6 (Figure 3D and 3E) with over 75% of ATF6 remaining after DTT treatment. Yet, processing of ATF6 constructs was not attenuated under *WT*-*L.p.* infection even in the absence of a functional S1P cleavage site (Figure 3D and 3E). Cumulatively, the data suggests *L.p.* infection stimulates the ATF6 pathway through means that circumvent the requirement of canonical pathway components.

**Figure 3.**
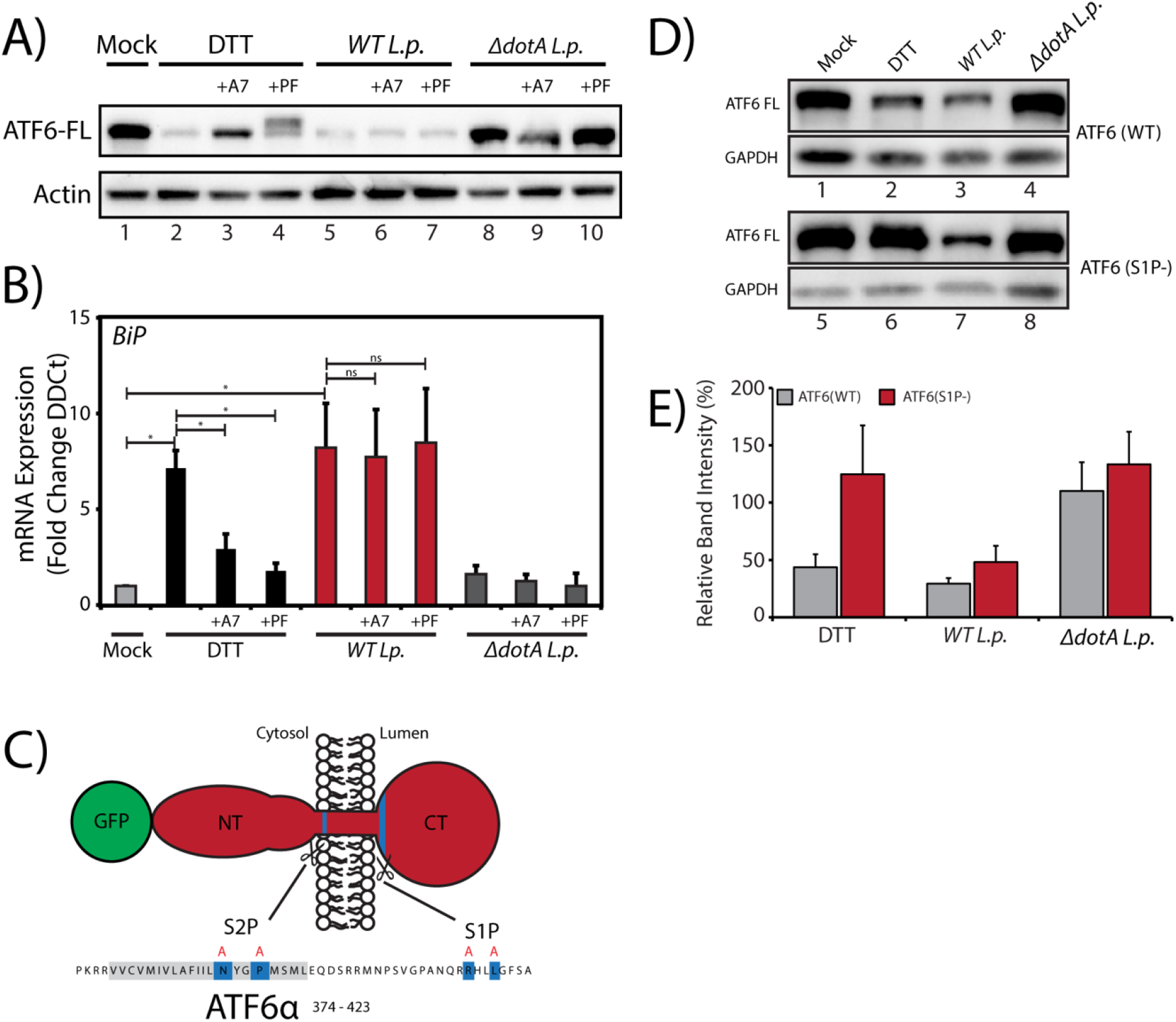
*L.p.* induces non-canonical activation of ATF6. (A, B) HEK293-FcγR cells were treated 6 μM Ceapin A7 or 1 mM PF-429242 for 1 hour prior to infection or DTT treatment. DTT treated samples were incubated with 1 mM DTT for (A) 1 hour or (B) 4 hour. Infected cells were incubated with *L. p.* (*WT* or *ΔdotA*) for 6 hours, then lysed and (A) analyzed by immunoblot analysis using anti-ATF6 and anti-Actin antibody, or (B) analyzed by RT-qPCR using primers against *BiP*. (C) Schematic of membrane bound N-terminal GFP-tagged ATF6 (red) with relevant features highlighting S1P and S2P cleavage site sequences (blue) and mutated residues for S1P and S2P cleavage site mutations (red letters). (D) HEK293-FcγR cells were transfected with GFP-ATF6(WT) or GFP-ATF6(R415A/R416A). Transfected cells were incubated with 1 mM DTT for 1 hour, or infected with *L. p.* (*WT* or *ΔdotA*) for 6 hours and analyzed by immunoblot using anti-GFP and anti-GAPDH antibodies. (E) Densitometry analysis showing the percentage of ATF6-FL signal remaining in treated cells relative to mock control cells.

### L. p. infection stimulates translocation of ATF6 to the nucleus

We next examined whether *L.p.* infection causes nuclear accumulation of ATF6. Using a previously validated N-terminal GFP fusion protein, GFP-ATF6, the canonical activation pathway as induced with DTT was monitored using live-cell time-lapse microscopy (Chen et al., 2002). HeLa cells stably expressing the Fc Gamma receptor (HeLa-FcγR) were co-transfected with GFP-ATF6 and Golgi marker GalT-RFP. DTT treatment caused a rapid translocation of ATF6 from the ER to the Golgi within 30 minutes, and ATF6 signal colocalized with the Golgi marker GalT-RFP (Figure 4A, Supplemental Movie S1). After 120 minutes of DTT treatment, the ATF6 signal was predominantly localized to the nucleus. We then monitored the ATF6 localization pattern during *L.p.* infection using JF-644-stained Halo-tagged *L.p.* strains (Figure 4B, Supplemental Movie S2) (Grimm, 2017). Analysis of time-lapse micrographs revealed an increase in nuclear signal intensity during *WT-L.p.* infection (Figure 4C, D). Strikingly, ATF6 activation during *L.p.* infection did not produce the robust Golgi/perinuclear recruitment as seen with DTT induction (Supplemental Movie S2). The ability of *L.p.* to activate ATF6 in a manner that bypasses a need for Golgi translocation further corroborates our S1P mutants and Ceapin A7 studies that showed that ATF6 processing does not occur at the Golgi during infection. As infection progresses, the LCV is remodeled from a plasma membrane derived vacuole to an ER-like compartment in a process that involves the recruitment of host ER proteins to the LCV and the disruption of ER-to-Golgi trafficking (Swanson, 1995, Tinley, 2001). When monitoring ATF6 localization, confocal microscopy revealed substantial recruitment of ATF6 to the LCV membrane at 6 hours post-infection with over 80% of LCVs marked positive for ATF6 (Figure 4E). Whereas a majority of LCVs were marked positive for ATF6 in WT-*L.p*., the recruitment required a functional Dot/Icm system as the *ΔdotA-L.p.* infected cells exhibited ATF6 recruitment to approximately 30% of LCVs. These data support the hypothesis that ATF6 is activated through a non-canonical mechanism.

**Figure 4.**
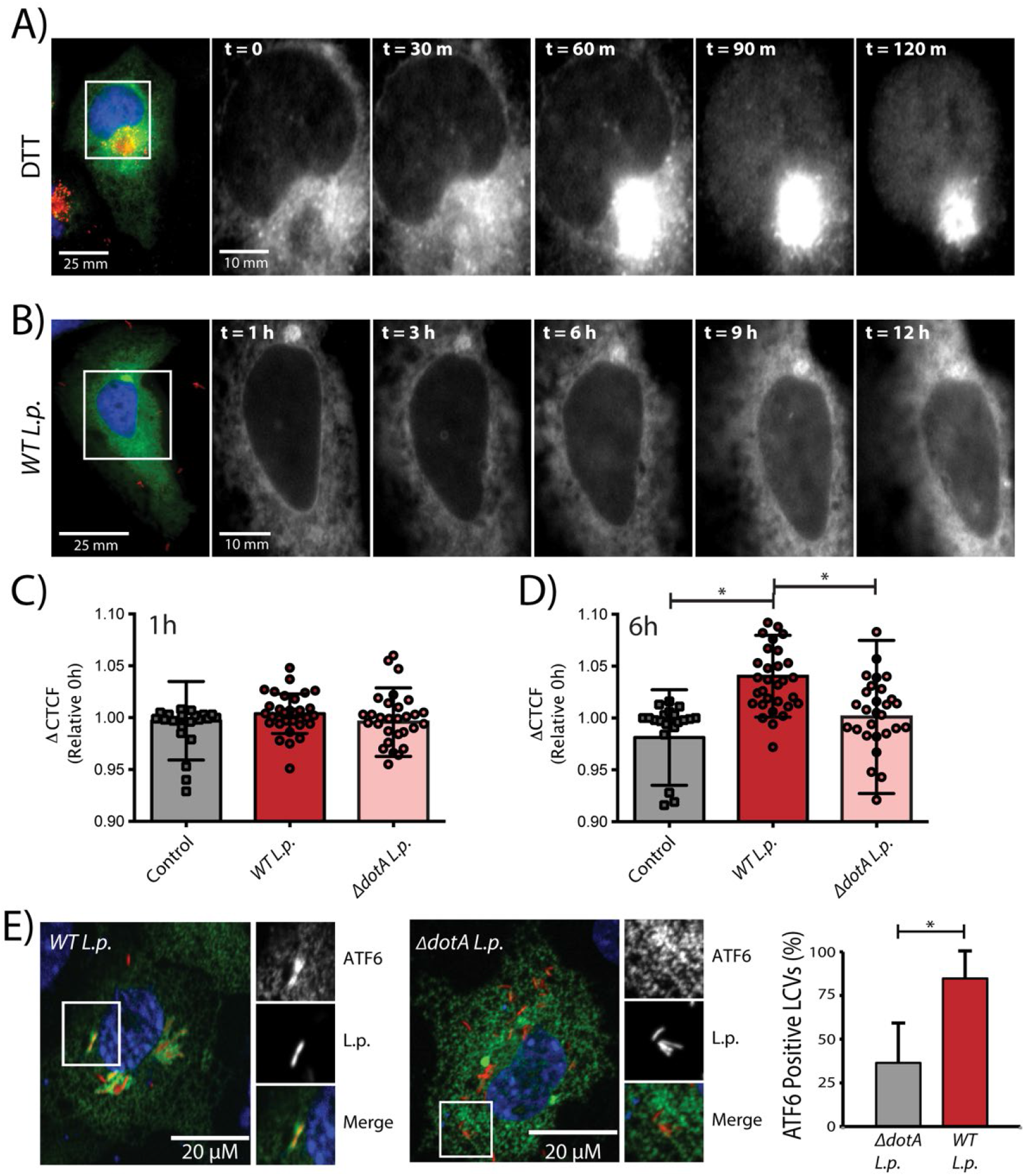
*L.p.* infection stimulates ATF6 nuclear recruitment. (A, B) Representative image stills from time-lapse wide-field fluorescent microscopy of HeLa-FcγR cells (A) transiently transfected with GalT-RFP and GFP-ATF6, then treated with 1 mM DTT or (B) transfected with GFP-ATF6, then infected with JF-646 stained *Halo-tag WT-L.p.* Live-cell epifluorescence image stills at timepoint 0 (t=0) (panel 1, scale bar = 25μm), with white box denoting location of inset. GFP-ATF6 signal (inset, panel 2-6, scale bar = 10 μm) (C, D) Quantitation of temporal changes in nuclear localized GFP-ATF6 relative corrected total cell fluorescence (CTCF) of ATF6 in single cells followed over a 9-hour period analyzed between 1 hour (C) and 6 hours. Signal intensities analyzed from of control (n = 25), *WT L.p. (*n = 30,) or Δ*dotA L.p.* (n = 30) at designated time points were normalized to signal at 30 min post infection. (E) Confocal micrographs of FcγRII and GFP-ATF6 transfected Cos7 cells infected with JF-646 stained *Halo-tag L. p. (WT or ΔdotA).* Values are the Mean ± SEM of three independent experiments in which at 50 LCVs were scored for each condition.

### ATF6 activation is strain and species specific

The data presented here highlights a T4SS-dependent activation strategy requiring the translocation of *Legionella* effector proteins. Therefore, we sought to identify effector proteins capable of inducing ATF6 activation by identifying *Legionella* strains that fail to efficiently process ATF6. Genomic analysis of over 30 *Legionella* strains and species revealed largely non-overlapping effector repertoires (Burstein, Amaro et al., 2016); therefore, we sought to use a comparative approach to test for ATF6 processing in different *Legionella* strains and species. Four *Legionella* species – *Legionella pneumophila*, *Legionella micdadei* (*L. mic)*, *Legionella wadsworthii* (*L. wad*), and *Legionella longbeachae* (*L. lon*) – were tested in addition to *Legionella pneumophila* strains – *Philadelphia* str. (*WT L.p.-Phila* or *ΔdotA L.p.-Phila*), *Paris* str. (*L.p.-Paris*), *Lens* str. (*L.p.-Lens*), and *Serogroup 6* str. (*L.p.-SG6*). The *Legionella* species and strains were used to infect HEK293-FcγR cells and endogenous ATF6 processing was monitored by immunoblot. While majority of the species and strains tested recapitulated the ATF6 processing seen with wild type *L.p.*, both *Legionella wadsworthii* and *Legionella pneumophila Paris* str. showed greatly reduced ATF6 processing (Figure 5A, B). To test whether the infection resulted in a functional *Legionella* containing vacuole (LCV), we next tested the recruitment of ubiquitin to the vacuole. Ubiquitination of the LCV has been seen as a hallmark of successful infection (Horenkamp, 2014). Whereas the *Legionella wadsworthii* strain failed to show ubiquitin recruitment, the *Paris* strain exhibited a robust recruitment of ubiquitin-modified substrates around the LCV in the HEK293-FcγR cells as observed by immunofluorescence (Figure 5C).

**Figure 5.**
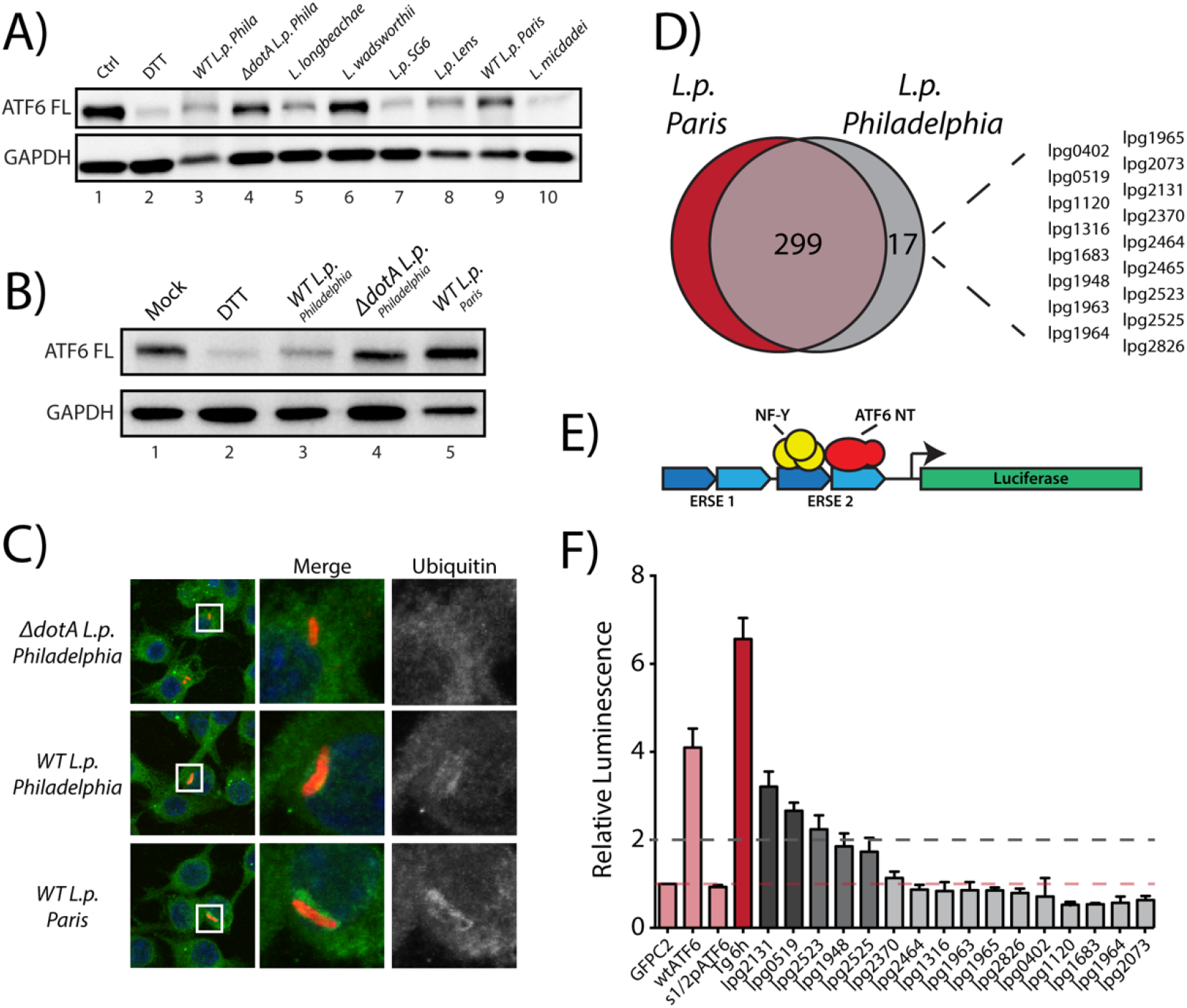
*L.p. Paris* strain does not induce ATF6 processing. (A, B) Immunoblots of HEK293-FcγR cells infected with *Legionella pneumophila* strains – *Philadelphia* str. (*WT L.p.-Phila* or *ΔdotA L.p.-Phila*), *Paris* str. (*L.p.-Paris*), *Lens* str. (*L.p.-Lens*), and *Serogroup 6* str. (*L.p.-SG-6*), and *Legionella micdadei* (*L. mic)*, *Legionella wadsworthii* (*L. wad*), and *Legionella longbeachae* (*L. lon*). (C) Immunofluorescence micrographs from *L.p. (WT, dotA, or Paris str.)* infected HEK293-FcγR cells stained with Hoechst (blue) and antibodies against Ubiquitin (green) and *Legionella* (red). (D) Identification of gene of orthologs of *L.p*. *Philadelphia* strain effectors through pairwise sequence alignments against *L.p. Paris* strain. Listed are *Philadelphia* strain effectors that are absent from the *Paris* strain. (E) Schematic representation of ERSE-luciferase construct stably expressed in a HEK-293T based cell line (HEK293T-ERSE-Luciferase). Two copies of the ER stress response element (ERSE, blue) are cloned upstream of a minimal promoter driving Luciferase (green) gene expression. Once cleaved, ATF6(NT) binds the ERSE sequence and stimulates Luciferase expression. (F) Primary screen data from ERSE-Luciferase reporter. HEK293T-ERSE-Luciferase cells were transiently transfected with Myc-tagged *Legionella* effectors or GFP-C2 vector, GFP-ATF6(WT), or GFP-ATF6(R415A/R416A, N390F/P393L) (S1P/S2P). Treatments were internally normalized to non-treated control cells. Fold induction of luciferase activity from transfected samples taken relative to empty control vector transfected cells is charted. Baseline luciferase activity from control cells (red line) and 2-fold induction cutoff (grey line) are indicated.

Comparative genomic analysis between the *Philadelphia* str. and *Paris* str. revealed 17 known *Philadelphia* str. effector proteins that were absent from *L.p. Paris* str. effector repertoire (Figure 5D). To identify effectors capable of activating the ATF6 pathway, a previously validated HEK293T cell line comprising of a stably integrated tandem ER stress response element with regulatory control over Luciferase expression (HEK-ERSE-Luc) (Gallagher, Garri et al., 2016) was used (Figure 5E). The 17 Myc-tagged effectors were transiently expressed into the HEK-ERSE-Luc cell line and screened for luciferase induction. UPR induction as caused by thapsigargin (Tg) treatment produced a greater than 6-fold induction of luciferase activity (Figure 5F, red). Control vectors expressing WT-ATF6 and S1P/S2P-ATF6-null were also evaluated in the luciferase cell line. As reported previously (Ye et al., 2000), overexpression on ATF6 led to an increased basal level of activation producing a 4-fold increase in luciferase activity over control vector, whereas the cleavage deficient ATF6 mutant did not lead to an increase in baseline luciferase activity. This data excludes the possibility that simply overexpressing an ER protein that trigger ATF6 expression. Our assay revealed that expression of 11 of the 17 effectors did not increase luciferase activity as compared to the control vector. While, *L.p. Philadelphia str.* effectors *lpg1948*, *lpg2523*, and *lpg2525* stimulated a 2-fold increase in luciferase activity, the effectors *lpg0519* and *lpg2131* produced a greater than 2-fold increase in luciferase activity when expressed individually (Figure 5F). Together, these results indicated that multiple effectors possess the capacity to induce the ATF6 pathway. Most of the effectors identified in this screen had little or no known function assigned to them.

### Lpg0519 localizes to the ER and activates ATF6

To experimentally validate the results from the screen, the ATF6 targeting effectors were transiently expressed in HEK293T cells and UPR activation was monitored by immunoblot. Upon Tg treatment, a reduction in endogenous ATF6 was observed by 2 hours (Figure 6A). By 4 hours of Tg treatment, the level of endogenous ATF6 was restored (Figure 6A). Downstream of ATF6, UPR induction through Tg treatment also resulted in elevated levels of BiP/GRP78 compared to untreated cells. More broadly, Tg treatment also stimulated PERK pathway activation (a transmembrane UPR sensor at the ER) as shown by elevated ATF4 levels after 4 hours. When compared to Tg treatment, cells transfected with the GFP control vector did not exhibit a reduction in ATF6 levels, and BiP and ATF4 levels were not increased compared to untreated cells (Figure 6A and 6B). However, cells transfected with C-terminally tagged GFP-Lpg0519 exhibited a dramatic reduction in endogenous ATF6 levels (Figure 6A). Further validating ATF6 activation, BiP levels were also elevated in GFP-Lpg0519 transfected cells (Figure 6B). However, the levels of ATF4 were not elevated in Lpg0519 transfected cells (Figure 6A). These results highlight the UPR pathway specificity by Lpg0519 in activating the ATF6 pathway without targeting the UPR more generally. To further characterize Lpg0519, we investigated its subcellular localization in mammalian cells. GFP-Lpg0519 was transiently transfected in U2OS cells, and then examined in live cells using confocal microscopy. Whereas the GFP control vector was uniformly distributed throughout the cell (Figure 6C), Lpg0519 co-localized with the ER-marker mCherry-ER-3 (Figure 6C). Altogether, these data suggest that Lpg0519 localizes to the ER and has the capacity to specifically induce the cytoprotective branch of UPR (ATF6), without affecting the apoptotic branch (PERK).

**Figure 6.**
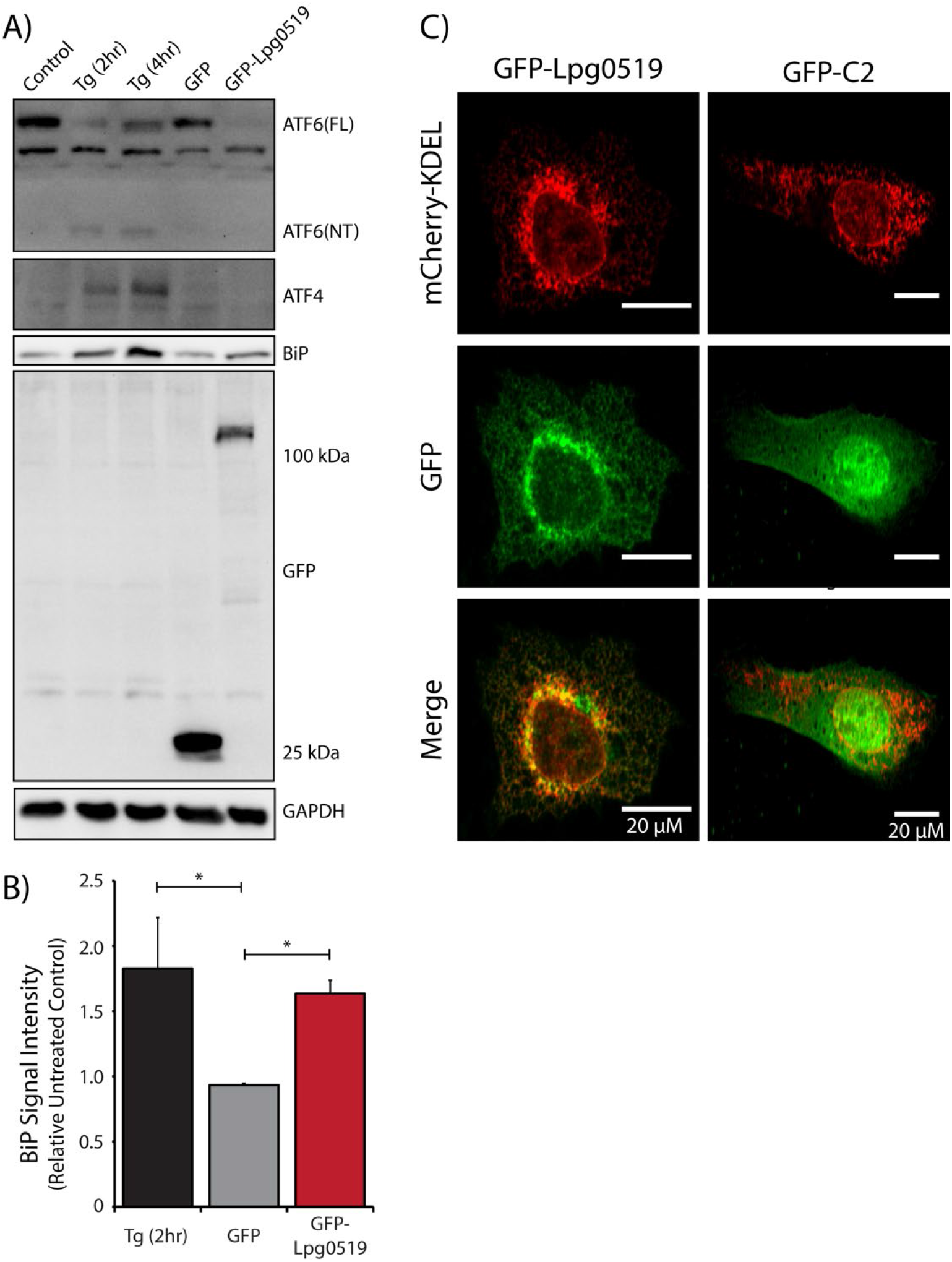
ER-localized Lpg0519 activates ATF6. (A) Immunoblot of HEK293T cells treated with thapsigargin (Tg) for 2 hours or 4 hours, or transiently transfected with GFP-C2 vector or GFP-Lpg0519. Antibodies against ATF6, ATF4, BiP, GFP, and GAPDH were used. (B) Quantification of replicate experiments (n=3) from (A) of BiP signal intensity relative non-treated control cells. Mean ± SEM, (*, p < 0.05). (C) Confocal micrographs from U2OS cells transfected with GFP-C2 vector (right, center) or GFP-Lpg0519 (left, center) and mCherry-Calreticulin (red, top).

## Discussion

The UPR represents a critical node for re-establishing ER homeostasis under perturbations caused by the accumulation of misfolded proteins. Importantly, the three branches of the UPR work synergistically to maximize the response, yet the modalities by which each sensor contributes to homeostatic restoration differ considerably (Ron & Walter, 2007). Of importance, the PERK and IRE1 pathways are also UPR components that integrate signals from pathogen associated molecular patterns that enable cross-talk between the UPR sensors and host cell innate immunity responses to defend again pathogen invasion (Martinon, 2010, Janssens, Pulendran et al., 2014, Smith, 2018). Therefore, the targeted inactivation of IRE1 and PERK mediated enhancement of cytokine production by *L.p.* likely contributes to pathogen survival. The ATF6 pathway has also been linked as a modulator of pro-inflammatory responses and ATF6 was shown to enhance NF-κB signaling in ER stressed macrophages (Rao, 2014). Paradoxically, multiple *L.p. effectors* have been identified that have the capability of activating this inflammatory response pathway (Losick, 2010, Ge, 2009).

As the ATF6 pathway is mainly associated with the production of chaperones, ERAD components and lipid synthesis enzymes, the pro-survival attributes associated with ATF6 activation make it an attractive target that might serve to benefit pathogen survival and replication. Especially, with respect to *L.p*., an attractive model to test in future studies would be to examine whether *L.p.* utilizes ATF6 for lipid synthesis to actively contribute to the growing membrane of the LCV during the course of the infection. Indeed, targeting of the ATF6 branch of the UPR has been utilized by protozoan, bacterial, and viral pathogens alike, and has been shown to contribute to intracellular replication in each system (Celli & Tsolis, 2015, Galluzzi et al., 2017, Jheng, Lau et al., 2010). Studies involving viruses have reported on a strategic activation of the UPR as a means to repurpose the autophagic response to benefit the virus where ATF6 knockdown reduced viral loads (Hou et al., 2019, Sharma, Bhattacharyya et al., 2017). However, this differs drastically from what is seen under *L.p.* infection, as the autophagy pathway is downregulated in *L.p.* infection (Choy, 2012). While bacteria have been shown to modulate UPR activity, it is striking how *L.p.* differentially modulates the different arms of the UPR by inhibiting IRE1, while activating ATF6 (Hempstead, 2015, Treacy-Abarca, 2015).

In this work, we aimed to further characterize the molecular mechanisms governing ATF6 processing in response to *L.p.* infection. ATF6 protein levels are tightly regulated through constitutive degradation via SEL1L/HRD1 dependent ERAD in conditions absent of ER stress and the impact of the constitutive degradation is revealed most strikingly under conditions of protein synthesis arrest (Fonseca, 2010, Horimoto, 2013). *L.p.*-induced translational inhibition occurs via host and pathogen mediated mechanisms and has been documented to have an impact on UPR induction under infection (Hempstead, 2015, Treacy-Abarca, 2015). Thus, we considered constitutive degradation of ATF6 to be the source for the reduction in protein levels during infection. Yet, we found impaired SEL1L-dependent ERAD and pharmacological inhibition of proteasomal degradation failed to stabilize ATF6 protein levels during infection (Figure 1D).

Though proteasomal degradation pathways are active, our data suggest that degradation of ATF6 is not a major contributor to the loss of ATF6 signal during *L.p.* infection. Furthermore, our data revealed that reduction in ATF6(FL) protein levels correlated with the appearance of an ATF6 cleavage fragment during wild type infections (Figure 2D). We considered the hypothesis that ATF6 is processed to form an active transcription factor capable of increasing the expression of target genes. Indeed, we found ATF6 regulated genes to be expressed under infection and silencing of ATF6 reduced target gene expression directly linking the sensor to downstream transcriptional activation (Figure 2A). Though many UPR genes are highly expressed during infection, the genes are not translated at the rates observed under ER stress (Hempstead, 2015, Treacy-Abarca, 2015).

We then determined host and pathogen components required for ATF6 activation under infection. To our surprise, the major host pathway components were dispensable for *L.p.* mediated activation. Blockage of ATF6 specific ER-to-Golgi translocation and inhibition of S1P/S2P proteolytic activity traditionally restrict ATF6 activation under ER stress, however, neither impacted *L.p.* induced pathway activation in terms of ATF6 processing and downstream gene activation (Figure 3A, 3B). Using time-lapse microscopy, we observed *L.p.* infection stimulated nuclear recruitment of ATF6 without the preceding Golgi translocation observed under ER stress induced activation (Supplemental Movie S2). Furthermore, ATF6 was detected around the LCV at 6 hours post infection suggesting a mechanism for direct recruitment to the pathogen vacuole (Figure 4E). More so, the capacity for wild type *L.p.* infection to induce processing of S1P and S2P site null mutants suggests a potentially unique site of cleavage under infection. However, the instability of the ATF6-NT cleavage fragment has so far hindered our efforts to the identify the novel cleavage site.

Our observations that ATF6 activation required a functional T4SS and the lack of a dependency for canonical host factors on ATF6 activation suggested one or more *L.p.* secreted effectors were involved. Using a phylogenetic approach, we identified the *L.p. Paris* strain lacks the capacity to cleave ATF6. In doing so, we were able to identify several effectors unique to the *L.p. Philadelphia* strain that, when expressed individually, increased expression of an ATF6 specific reporter (Figure 5F). Though the *L.p. Paris* strain lacks effectors shown to cleave ATF6, we do not dismiss the potential for the *Paris* strain to also contain negative regulators of ATF6 cleavage which could contribute to the observed reduction in ATF6 activation. Taken together, our results are the first example of a bacterial pathogen that can activate the ATF6 pathway independently of host pathway factors. Future work on these newly identified effectors might elucidate novel mechanisms of ATF6 activation and has the potential to open up new paradigms by which intracellular pathogens manipulate the cytoprotective UPR sensor ATF6.

## Materials and Methods

### Bacterial strains

All *Legionella* strains were gifts from Craig Roy’s laboratory at Yale University. *Legionella* strains used in this study were routinely cultivated on Charcoal Yeast Extract (CYE) agar. The *ΔdotA*, *ΔsidC-sdcA*, and IPTG-inducible Halo-expressing strains were derived from the parental *Lp01* strain. The *Δ2,3,4,6,7*, *Δ2,3,6,7*, *Δ2,3,6 L.p.* strains (O’Connor et al., 2011), and *Δ7-translation L. pneumophila* strain (Barry, 2013) was a gift from Russell Vance’s lab and are thymidine auxotrophs derived from the *Legionella pneumophila serogroup* 1 Lp02 (Berger and Isberg, 1993). *Legionella pneumophila Paris B1* strain was purchased from ATCC (ATCC 700833). Chloramphenicol (10 ug/mL), IPTG (0.1 mM), and thymidine (100 ug/mL) were added to CYE agar plates as needed. *L.p.* were harvested from 2-day heavy patches and used to infect cells.

### Cell culture

HEK-293 FcγRII and HeLa FcγRII cells were obtained from Craig Roy’s laboratory at Yale University. All cells were cultured in DMEM (Life Technologies) supplemented with 10% fetal bovine serum (FBS) at 37 °C and 5% CO_2_. RAW 264.7 macrophages were cultured in Roswell Park Memorial Institute media (RPMI) (Corning) supplemented with 10% FBS at 37 °C and 5% CO_2_. Cells were placed in poly-L-lysine–treated plates and grown to 90% confluency. Drug treatments were performed at final concentration, 200 nM Tg (Enzo Life Sciences, Farmingdale, NY), 1 mM DTT (Research Products International (RPI), Mt Prospect, IL), 300 μM 4-(2-aminoethyl)benzenesulfonyl fluoride (AEBSF) (Sigma-Aldrich, St. Louis, MO), 1 μM Pf-429242 (Sigma-Aldrich), 20 μM MG-132 (Enzo Life Sciences), 25 μM Cycloheximide (Sigma-Aldrich), 10 μM Ceapin A7 (Gallagher et al., 2016). Ceapin A7 was provided as a gift from the Peter Walter laboratory at UC San Francisco.

For transient transfections, cells were grown to 70% confluency and transfected with 2 μg of plasmid for 60 mm and 35 mm dishes, or 1 μg per well for 24-well plates using JetPRIME (Polyplus-transfection) according to the manufacturer’s instructions. Cells were incubated with transfection reagent for 4 hours, then media replaced with fresh DMEM supplemented with 10% FBS. For siRNA transfections, cells were grown to 30-50% confluency and transfected using Oligofectamine (Thermo Fisher Scientific, Waltham, MA) according to the manufactures protocol. Cells were grown for 72 hours prior to application of treatment conditions. The following siRNA oligos used were purchased from Sigma-Aldrich: SEL1L-TTAACTTGAACTCCTCTCCCATAGA, Scramble-GCATACTCAACTACTTCGCATACTT; ATF6-GAACAGGGCTCAAATTCTC, Scramble-GCTAGTGCACAAGTACCTA.

### Infection

Cells were infected at a multiplicity of infection (MOI) of 100 or 5. If cells required opsonization, *Legionella* polyclonal antibody (Invitrogen, Cat: PA1-7227) was used at 1:2,000 and incubated for 20 min at room temperature. Immediately after infection, cells were centrifuged for 5 min at 1,000 rpm. After centrifugation, cells were left at 37 °C for an additional 60 min. After 1 hour, cells were washed with 1x PBS to remove extracellular bacteria. DMEM supplemented with 10% FBS or the same media supplemented with treatment reagent was replaced after the washes in PBS. Infected cells were harvested at the designated times.

### Quantitative RT–PCR

HEK-293 FCγ cell mRNA was harvested and isolated using Direct-zol™ RNA Miniprep Plus (Zymo Research, Irvine, CA) according to the manufactures protocol. cDNA synthesis was performed using QuantiTect Reverse Transcription Kit (Qiagen, Hilden, Germany) and cDNA reactions were primed with poly dT. Relative Quantitative PCR was performed using iTaq Universal SYBR Green Supermix (Bio-Rad, Hercules, CA). Endogenous GAPDH mRNA was used for normalization. Uninfected and untreated HEK-293 FCγRII cells were used as the endogenous control for each qPCR analysis. The following RT–qPCR primers were used: BiP (Human) forward-CATCACGCCGTCCTATGTCG, reverse-CGTCAAAGACCGTGTTCTCG; HERPUD1 (Human) forward-AACGGCATGTTTTGCATCTG, reverse-GGGGAAGAAAGGTTCCGAAG; Sel1l (Human) forward-AAACCAGCTTTGACCGCCAT, reverse-GTCATAGGTTGTAGCACACCAC; Hyou1 (Human) forward-GAGGAGGCGAGTCTGTTGG, reverse-GCACTCCAGGTTTGACAATGG; ATF6 (Human) forward-AGAGAAGCCTGTCACTGGTC, reverse-TAATCGACTGCTGCTTTGCC

### Immunoblot analysis

Mammalian cells were lysed in Radioimmunoprecipitation assay buffer (RIPA) buffer with the addition of protease (Roche cOmplete), and phosphatase inhibitors (GB Sciences). Protein levels of lysates were determined using the BioRad DC/RC assay. Equal amounts of protein lysate were boiled with SDS load buffer, and equal amounts of protein were loaded. Immunoblotting was performed with the following antibodies: GAPDH (Proteintech, Cat: 60004- 1-lg), ATF6 (Proteintech, Cat: 60004-1-lg), beta-Actin (Proteintech, Cat: 20536-1-AP), alpha-Tubulin (Proteintech, Cat: 66031-1-Ig), Sel1L (Abcam, Cat: ab78298), GFP tag (Proteintech, Cat: 66002-1-Ig), BiP (Proteintech, Cat: 11587-1-AP), ATF4 (Cell Signaling Technology, Cat: 11815S).

### Immunofluorescence Microscopy

Cells were plated on 12 mm glass coverslips in 24-well plates. After 24 hours, cells were fixed, treated with drugs, or infected with *L.p.* as needed. For ubiquitin recruitment assay RAW cells were infected with *L.p.* at MOI = 5. 1 hour after infection, cells were washed twice with PBS to remove extracellular bacteria and incubated for 2 hours more. For co-localization assays, U2OS cells were co-transfected with pcDNA-FcγRII (Arasaki, 2010) and GFP-ATF6α, then infected with *L.p.* at MOI = 10 and infected for 1, 4, or 8 hours. For 4- and 8-hour time points, cells were washed with PBS after 1 hour to remove extracellular bacteria. Coverslips were mounted with ProLong Diamond Antifade Mountant (Thermo Fisher Scientific) incubated at 37 C for 10 minutes, then imaged directly. For immunofluorescence, coverslips were washed with cold 1x PBS, fixed with 4% paraformaldehyde in PBS for 10 min at room temperature, permeabilized in 0.1% saponin in PBS, and blocked in 3% bovine serum albumin (BSA) in PBS, and then incubated with the appropriate primary and secondary antibodies diluted in 3% BSA. Nuclei were stained with Hoechst 33342 dye for 10 min before mounting on microscope slides.

Coverslips were imaged using an inverted Nikon Eclipse Ti-E spinning disk confocal microscope equipped with a Prime 95B 25mm CMOS camera (Photometrics) camera. Antibodies used were Ubiquitin (Millipore, Cat: ST1200-100UG), and Secondary Antibody, Alexa Fluor 546 or Alexa Fluor 488 (Thermo Fisher Scientific).

### Live Cell Microscopy

HeLa FcγRII cells expressing GFP-ATF6 were plated on 35 mm poly-lysine-coated imaging dishes (Cellvis, Mountain View, CA). Cells were infected at MOI = 5 with *WT*- or *ΔdotA L.p.* that were previously stained with HaloTag-Janelia Fluor 646 conjugates (Grimm et al., 2017). For staining, briefly, *L.p.* maintaining a HaloTag-expressing plasmid were harvested from a 2-day heavy patch and incubated at liquid culture overnight with 0.1 mM IPTG. Liquid culture at OD = 3.0 was then pelleted and resuspended in 5 mM Janelia Fluor 646 HaloTag ligand (JF646) in order to facilitate HaloTag-JF646 conjugation. After 15 min incubation with ligand in the dark, *L. pneumophila* were washed 1x with water. Stained bacteria were resuspended in 2 mL of DMEM lacking phenol red (Gibco) and used for infection of cells. Imaging of cells took place in a controlled chamber maintaining 37C with 5% CO_2_. A random selection of cells was imaged at 60X magnification at 5- or 10-minute intervals for 12 hours using a Nikon Eclipse Ti2 microscope with a Nikon DS-Qi2 camera. Cells that died or lost focus over the time course were omitted from analysis.

### Luciferase assay

ONE-Glo Luciferase assay system was purchased from Promega. HEK-293T ERSE-Luciferase cells, provided as a gift from the Peter Walter laboratory at UC San Francisco and described previously (Gallagher et al. 2016), were seeded onto 6-well dishes at 70% confluency. Cells were transfected with Myc-tagged *Legionella* effectors as previously described (Barry, 2013) for 24 hours. Cells were suspended in 1mL DMEM and 50 uL aliquots were transferred to assay plate (Falcon, Cat:353296) in triplicate. One-Glo Luciferase reagent was pre-equilibrated to room temperature and added at equal volume into each well. Assay plate measurements were performed on Tecan Saphire^2^ with 1 s exposure. Background signal was subtracted from untreated untransfected cells. Treatment conditions were normalized to control Myc-tag expressing cells.

### Quantification Statistical analysis

GraphPad Prism 6 software was used for statistical analysis. Where statistical analysis was performed an unpaired student’s t-test was performed using three biological replicates. Statistical significance (*) was determined as a p-value < 0.05. Image analysis was performed using ImageJ.

## Acknowledgements

We thank Dr. Philipp Schlaermann for doing preliminary studies on ATF6. We thank Peter Walter for scientific advice and for very generously providing us with the ERSE cell-line and Ceapin A7 inhibitor. We thank members of the Mukherjee lab for critical reading of the manuscript. S.M. is supported by the National Institutes of Health RO1 grant AI118974 and an award from the Pew Charitable Trust (A129837).

## Author contributions

Conceptualization, S.M., N.U.I., Methodology, S.M and N.U.I.; Data Analysis, S.M. and N.U.I.; Investigation, N.U.I., Writing – Review & Editing, N.U.I., and S.M.; Funding Acquisition, S.M.; Resources, S.M.; Supervision, S.M.

---

Supplementary Movie S1 Time-lapse widefield microscopy movie of DTT treated HeLa-FcγRII cells expressing GFP-ATF6 (green) and GalT-RFP (red) and stained with Hoechst solution (blue). DTT was treated to cells at t=0. Still frame images were taken every 5 minutes over 2.5 hours.

Supplementary Movie S2 Time-lapse widefield microscopy movie of HeLa-FcγRII cells expressing GFP-ATF6 (green) and stained with Hoechst solution (blue). Cells were infected with JF-646 stained *Halo-tag WT-L.p.* Infected cells were imaged directly after centrifugation with no more than 30 minutes elapsed time between centrifugation and image acquisition at t=0. Still frame images were taken every 10 minutes over 12 hours.

